# Ecological monitoring using Collembola metabarcoding with extremely low bycatch amplification

**DOI:** 10.1101/2023.05.23.541478

**Authors:** Pedro M. Pedro, Laury Cullen, Fabiana Prado, Alexandre Uezu, Ross Piper, Christiana M.A. Faria, Christoph Knogge, Maria Tereza Pepe Razzolini, Marcela B. Paiva, Milena Dropa, Miriam Silva, Tatiane Cristina Rech, Thomas Püttker

## Abstract

Collembola are used widely to monitor soil health and functional parameters. Recent developments in high throughput sequencing (especially metabarcoding) have substantially increased their potential for these ends. Collembola are especially amenable to metabarcoding because of their small size, high abundance, and ubiquity in most habitat types. However, most Collembola sampling protocols collect a substantial and highly varied bycatch that can be a considerable impediment to metabarcoding, especially because of data lost to non-target species. We designed a primer set amplifying the D2 expansion segment of ribosomal DNA that is highly conserved across Collembola and successfully excludes from amplification nearly all other invertebrate taxa. We tested the diagnostic power of the primer set by clearly distinguishing Collembola communities between forest sites with differing habitat qualities in São Paulo State, Brazil. The oligos successfully amplified targets from all Collembola orders previously encountered in the sampling locations, with no non-target amplification, and also excluded the closely related Protura and Diplura. Alpha diversity (OTU count) and phylogenetic diversity was significantly higher in high quality habitats. Moreover, the beta diversity indices successfully differentiated high and low-quality habitats. This new addition to the biomonitoring toolbox greatly increases the accessibility of Collembola metabarcoding for various types of habitat assessments.

## Introduction

Collembola (springtails) are important invertebrate indicators of soil ecological parameters (Bispo et al., 2009) and are organisms particularly amenable to biomonitoring initiatives (Breure et al., 2003; Fiera, 2009; Filho et al., 2016; Zeppelini et al., 2009). The taxon is comprised of species highly adapted to specific local conditions and their ubiquity in most soil environments enables reliable spatial and temporal comparisons across a broad spectrum of habitats (Ponge, 1993).

Collembola are also important for *in vitro* toxicology assessments; they are especially utilized to detect harmful levels of heavy metals (Fountain and Hopkin, 2001; Lors et al., 2006).

Collembola are a statistically appealing taxon because diversity calculations can be based on high sample sizes per collection, whilst often maintaining a low sampling effort. In tropical soils, for example, Collembola can comprise over 60,000 individuals per m^2^ and are often represented by more than 100 species (Basset et al., 2022; Culik et al., 2002).

The taxonomic bottleneck currently limits Collembolan’s full potential as an efficient soil biological indicator. Morphological identification is particularly difficult because of the small size of most taxa and a dearth of diagnostic morphological traits resulting in high occurrence of cryptic species (Porco et al., 2012; Sun et al., 2017). However, a growing DNA barcode reference library has largely made this “impediment” moot. Genetic markers can now provide effective identification of many species, regardless of their life stage and/or specimen integrity (Beng et al., 2016; Eaton et al., 2017).

Recently, metabarcoding pipelines have greatly broadened springtails’ potential as biological indicators (Saitoh et al., 2016). Metazoan *metabarcoding* has built upon the advantages of single-specimen *barcoding* to simultaneously identify potentially thousands of individuals per sample (Yu et al. 2012, Ji et al., 2013). However, the technique has one significant drawback that can result in substantial analytical inefficiencies, particularly when using non-selective field sampling protocols: a significant portion of the DNA sequences resulting from bulk samples can comprise non-target DNA, which occupies dead space in sequencing flow cells and increases costs. To circumvent this issue, researchers can use primers targeting focal taxa, such that little or no *a priori* hand-sorting is required prior to DNA extraction (Brown et al., 2016; Ficetola et al., 2008, Pedro et al., 2020)

The concentration of Collembola DNA can often be overwhelmed by DNA from non-target bycatch. However, the most popular sampling protocols of Collembola (e.g. Berlese-Tullgren funnel, pitfall traps) are nonselective and thus are highly susceptible to collect substantial non-target bycatch of other invertebrates (Querner and Bruckner, 2010). The time investment needed to process these samples is often infeasible for practical biomonitoring.

Here we present a primer set designed to amplify an approximately 440-bp fragment of the Collembola D2 expansion segment of the 28S operon without amplifying significant non-Collembola bycatch (validated *in silico* and *in vitr*o). The primers’ exclusivity to Collembola was benchmarked by sequencing the entire contents of field-set pitfall traps, with minimal or no sorting. We assessed if this primer set, and metabarcoding analytical pipeline could be used in normal sampling protocols to test for differences in community composition between mature forests and incipient reforestation plots.

## Methods and Materials

### Primer design

Ribosomal loci are used in metabarcoding initiatives because they are taxonomically informative and, because of their non-protein-coding nature, often allow for the design of primers fully conserved to the target region (i.e., with no degeneracy at third codon positions; Burki et al., 2021; Semmouri et al., 2021). There is expected to be minimal PCR-bias when using these types of primers and rDNA is therefore an attractive option when estimates of relative proportions of taxa are needed (e.g., Pedro et al., 2020). Here, we target the D2 domain of the 28S operon of nuclear rDNA, which has been extensively used in species diagnosis, both pre- and post-metabarcoding (Campbell et al., 1994; Dodd et al., 2000; Pedro et al., 2021).

We initially downloaded all available GenBank accessions with keyword “28S” for Collembola and non-Collembola arthropods that contained either of the conserved D2 flanks using default MegaBLAST parameters (as per GenBank accession JX261730 for *Heteromurus* sp.). Sequence hits flanking the forward annealing region totalled 1,492 for Collembola and 42,129 for non-Collembolan arthropods. Reverse sequences were 562 and 38,200, respectively. Non-arthropods were not considered, as their D2 flanks were considerably diverged from the target ingroup and thus unlikely to compete with our target Collembola templates.

We evaluated only BLAST results with full binomial identification (= genus + species) and filtered results to exclude hits where putative priming sites were less than 20 nucleotides from the sequence termini (in order to omit those unwittingly submitted to GenBank with the original amplification oligo sequences still included). The filtered Collembola D2 sequences comprised 13 families and represented all four currently recognized Collembola orders.

Forward and reverse primers were designed that amplified the polymorphic region of D2 and were conserved among all available filtered Collembola GenBank sequences (576 and 166, respectively) with little or no degeneracy. These were designed so that the non-Collembola sequences did not possess the matching 3’ nucleotide in either primer. The result of these comparisons was *Collembola-F* (5’-AGAGAGTTMAAWAGTACGTGAAACCT-3’) and *Collembola-R* (5’-TGTTTCAAGACGGGACAGGC-3’).

We graphically confirmed the binding site variation and taxonomic resolution of the *Collembola-F* and *Collembola-R* primers designed above against all Collembola GenBank entries possessing both forward and reverse priming locations using the *ecopcr* module of Obitools (v1.2.13) (Boyer et al., 2016). We undertook an analogous comparative evaluation using two primer sets previously used in Collembola metabarcoding, 16S and CO1 (Saitoh et al., 2016), to assess the appropriateness of all currently available oligos. Our parameters for *ecoprc* allowed for a maximum of four mismatches in either primer and sequences had to possess priming sites at least 20 nucleotides from their termini. Expected *ecopcr* amplicon length in arthropods for D2 was 200-600 bp, for CO1 was 260-280 bp and for 16S 300-600 bp.

We also tested the fidelity of D2, 16S and CO1 primers to three non-Collembola taxa commonly found during pitfall and funnel sampling methods (based on our previous experience with these protocols): Acariformes, Coleoptera and Hymenoptera (principally ants). Here, we sought to assess each primer set’s cross-amplification in these non-target groups.

The resulting *ecopcr* output was analysed and graphed in R (R Development Core Team 3.0.1, 2013) using the ROBITools package (http://metabarcoding.org/obitools).

### Pitfall sampling

In order to test the applicability of the D2 primer sets in analyzing richness and composition of Collembola communities, we used them on invertebrate samples obtained by pitfall sampling in two locations in São Paulo State, Brazil. The first location, in Ubarana municipality (approximately 21°14’09” S, 49°43’12” W), consisted of a forest remnant adjacent to a hydroelectric reservoir, within which invertebrates were sampled at 13 sampling points. The second location was located near Nazaré Paulista (23°12’48” S, 46°21’58” W) where 15 points were sampled. In both locations, sampling points were distributed within forest classified as high-quality as well as low-quality (Supplemental Data 1). The latter category comprised areas of reforestation initiatives in initial stages of succession.

### DNA extraction, PCR and sequencing

The contents of pitfall traps were returned to the laboratory and stored at −20°C for not more than 1 week prior to DNA extraction. The 50-ml tubes used as pitfalls contained substantial amounts of bycatch, dominated by soil mites, small beetles and ants. We decanted the contents of the 50-ml tubes into weighing boats and then removed bycatch that was cumbersome to the downstream protocol, such as very large-carapaced insects and sources of PCR inhibitors (twigs, leaves, soil), but no further sorting was done, as taxon-specific primers were used. Weighing boat contents were transferred to 2-ml Eppendorf tubes, dried overnight on silica gel, and macerated using a Savant FastPrep lysis mill at maximum speed for 20-s using 1-mm ceramic beads (when necessary, large samples were divided into multiple tubes that were re-pooled following maceration).

A subsample of the maceration product was then submitted to DNA extraction using a DNeasy Blood & Tissue Kit (Qiagen, Valencia, CA) following manufacturer’s instructions for insects.

The working stock DNA from each sample extraction was diluted to 100ng/µl. Nested PCR reactions were performed using the Collembola-specific D2 primer set described above. Samples from different collection sites were tagged with multiplex identifiers (MIDs) to allow combined 454 FLX sequencing. In summary: a first PCR was done using the forward primer *Collembola-F_adF* (5’-<u>GGCCACGCGTCGACTAGTAC</u> AGAGAGTTMAAWAGTACGTGAAACCT-3’), where the underlined portion is an adaptor overhang used to decrease the cost of multiplexing PCR primers. The reverse primer for the first PCR was *Collembola-R* (5’-TGTTTCAAGACGGGACAGGC-3’).

The product from the initial reaction was diluted 10x, purified and submitted to a second PCR using the forward primer *454A-MID-adF* (5’-*CGTATCGCCTCCCTCGCGCCATCAG* NNNNNN <u>GGCCACGCGTCGACTAGTAC</u>-3’; where Ns represent a 6-bp barcode, the forward 454 fusion primer is italicized, and the adaptor overhang sequence is underlined) and the reverse *Collembola-R_454B* (5’-*CTATGCGCCTTGCCAGCCCGCTCAG* TGTTTCAAGACGGGACAGGC-3’), where the 454 fusion reverse primer is italicized.

Clean products (QIAquick PCR Purification Kit) were sequenced in the forward direction on 1/8 plate of a 454 Life Sciences Genome Sequencer FLX machine (Roche, Branford, CT) using the Macrogen facilities (South Korea).

### Sequence processing

We used MOTHUR v.1.36.1 (Schloss et al., 2009) to filter NGS sequences with a minimum average quality score of 25 and minimum length, after trimming of primer sequences, of 150-bp. We allowed for no nucleotide differences in the barcode region of the oligo and four differences in the priming region. Clustering of reads into OTUs was done as described in the USEARCH 454 SOP (http://drive5.com/usearch/manual8.1/upp_454.html;Edgar, 2010), which also removes chimeras based on de novo detection. OTUs represented by nine or fewer reads were removed from all subsequent analyses. A read-clustering threshold of 3% was adopted to bin OTUs. We assigned OTUs to taxonomy using a database created from all available Collembola GenBank entries using the RDP classifier (Wang et al., 2007).

### Alpha and beta diversity estimates

In order to evaluate the influence of habitat quality on Collembola richness, we estimated the number of operational taxonomic units (OTUs; MOTHUR command *summary*.*single*) and the scaled phylogenetic diversity (command *phylo*.*diversity*) at each sampling location. For each of the two response variables, we ran a model selection based on a candidate model set including “forest quality” and “sampling month” as fixed factors, and “sampling region” as a random factor.

To test for differences in community composition, we calculated pairwise dissimilarity between all sampling points relying on i) the Jaccard index (occurrence-based index) and ii) the Bray-Curtis dissimilarity index (abundance-based), and used Non-metric multidimensional scaling (NMDS), as well as Permutational multivariate analysis of variance (PERMANOVA; Anderson, 2006) (see Supplemental Data 2 for details on statistical analyses). All analyses were carried out in R using lme4 (Bates et al., 2015) and vegan package (Oksanen et al. 2022).

## Results

### Primer design

The GenBank sequences employed in the initial primer design were used to create the *ecopcr* libraries. For Collembola, 278 D2 sequences conformed to the *ecopcr* parameters, i.e., possessed both forward and reverse primers that were at least 20-bp away from sequence end. Another 26,468 non-Collembola arthropod D2 sequences were retrieved that matched the parameters above.

These represented all four Collembola orders and their priming sites were nearly universally conserved except for two degenerate positions in the forward primer (Figure 1). Although we were compelled to include two degenerate positions in the primer *Collembola-F*, these were 15 and 18 nucleotides away from the 3’-end, locations that generally result in little PCR bias (Kwok et al., 1990). A GenBank entry for *Archisotoma besselsi* (acc. No: JN981045) was the only Collembola D2 sequences to have a 3’ mismatch to the *Collembola-F* primer. We cannot discount that this is indeed a D2 polymorphism in class Collembola, although this nucleotide is not mismatched in any other representative of family Isotomidae.

**Figure 1:**
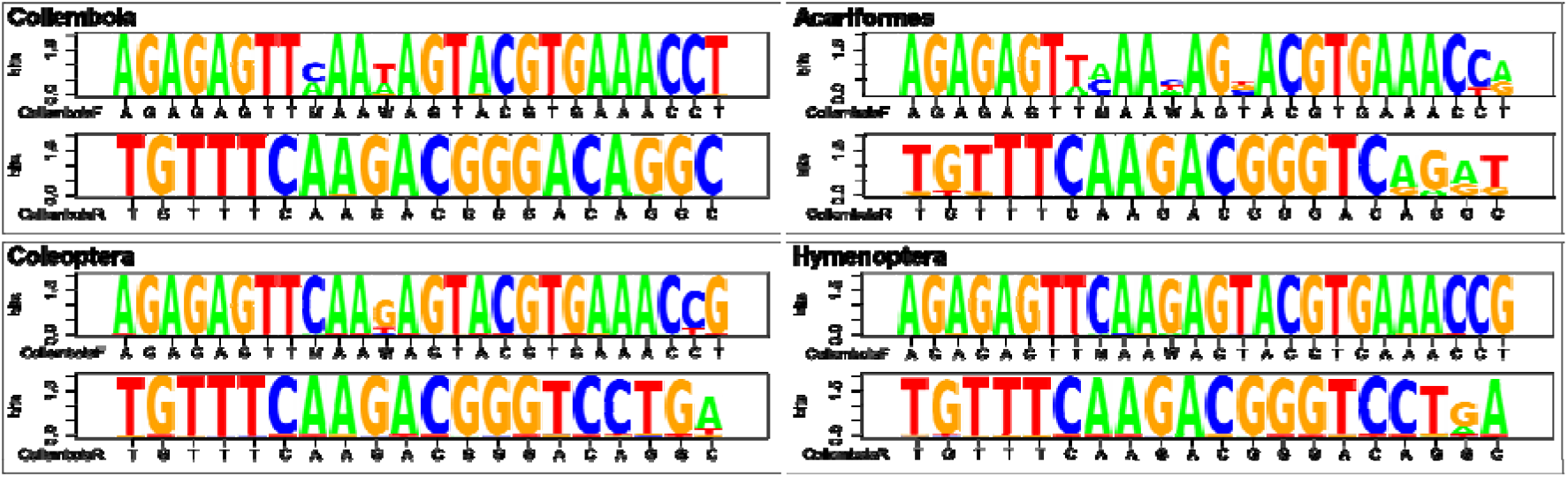
Sequence logos of the D2 primers designed herein (*Collembola-F* and *Collembola-R*) assessed for their complementarity to Collembola and three additional taxa commonly found during pitfall sampling protocols: Acariformes, Coleoptera and Hymenoptera.

Our primer design strategy for Collembola sought to maximally exclude non-Collembola amplification by having 3’ nucleotides that mismatched non-Collembola. There were few GenBank sequences available for the two other Entognatha, Protura and Diplura (less than 25 entries for each). However, in those sequences that possessed the primer sites, the 3’ nucleotides of both the forward and reverse primers mismatched that of the D2 primers designed herein, suggesting that, even in the presence of these closely related taxa, amplification should occur in only Collembola targets.

The 3’-T in *Collembola-F* and 3’-C in *Collembola-R* were rarely found in any non-Collembola arthropod sequences (in 0.06 % and 4.6 % of those conforming to the *ecopcr* parameters, respectively), and in these instances the two primers never occurred together in an individual sequence, thus precluding PCR. Moreover, within the three focal taxa assessed in greater detail, only 0.096 % Coleoptera sequences (n=6,829) possessed the 3’ match in the forward primer and 1 % had the 3’ nucleotide in the reverse priming location. In GenBank entries for Hymenoptera (n=5,224), 1.6 % had a 3’ nucleotide in the reverse primer designed herein (Figure 1). Neither of the 3’ nucleotides matched GenBank entries for Acariformes (from a total of 78 sequences), thus removing from possibility amplification of another very common soil invertebrate that is nearly always co-sampled with Collembola.

Conversely to the D2 primers, substantial primer conservation (both generally and at the 3’ nucleotide) among all Metazoa was seen in the two primer sets previously used in Collembola metabarcoding (Saitoh et al., 2016). For example, the 16S priming positions were relatively well conserved amongst the Collembola sequences (thereby minimizing primer bias), but also were not substantially different at their 3’ end from the other taxa evaluated (Supplemental Data 2, Figure S1). Likewise, the CO1 primers were equally similar to Collembola and to non-Collembola targets (Supplemental Data 2, Figure S2). Moreover, the CO1 primers were polymorphic at silent substitutions at each third codon positions, even those near the sensitive 3’ end. This may increase primer bias (Arnheim and Erlich, 1992) and, consequently, limit comprehensive estimates of relative abundances of PCR targets in bulk samples.

Amplicon length (including primers) varied little between the Collembola species assessed by *ecopcr* (440 bp SD=40) and the percentage GC composition was constant 48% (SD=4%). The taxonomic resolution of the D2 primers was relatively similar to both the COI and 16S sets previously used by Saitoh (2016b). In comparisons of only those species that were shared between the two markers, D2 (36 sequences) and 16S (42 sequences) both distinguished 100% of species. The COI primer set has marginally better resolution in 718 GenBank sequences than D2 (89 sequences): 96.9% versus 95.4%.

### Pitfall trapping

Some sampling sites were either vandalized or destroyed by wildlife (one in Ubarana and six in Nazaré Paulista) and therefore omitted from downstream analyses. Final sampling site totals were n= 9 in high-quality and n= 4 in low-quality forests in Ubarana; n=4 in high quality and n= 11 in low quality forests in Nazaré Paulista.

Although all pitfall traps contained much more bycatch DNA (from Diptera, Lepidoptera, Coleoptera and Hymenoptera (primarily ants)) than Collembola tissue, we nonetheless limited sorting effort to the rare cases of twigs, soil or very large beetles present amongst the catch.

### PCR and sequencing results

The first of the nested PCRs did not produce primer dimers or non-specific bands (Supplemental Data 2, Figure S3). This substantially simplified the laboratory pipeline, as a clean-up step was not needed before the second, indexing PCR.

Following sequence filtering, 35 unique OTUs were derived from the two sampling locations, averaging ∼460-bp. RDP results assigned all of these to class Collembola (100% bootstrap; see Supplemental Data 3), indicating no cross-amplification of bycatch DNA. All OTUs were assigned to genera at RDP 80% threshold or above, except OTUs 36, 27, and 8, which had support values of 72, 72, and 57 % respectively. The RDP results spanned ten of the 19 Collembola families currently represented in Brazil (Abrantes et al., 2010). Of the families previously recorded in Sao Paulo State, only Arrhopalitidae, Brachystomellidae and Tullbergiidae were undetected in our samples. Resulting sequences were assigned to all four Collembola orders except Neelipleona, which has not previously been documented in Sao Paulo State.

Collembola OTUs were highly specific to either of the two localities, potentially indicating a distributional or ecological effect on diversity in these two Atlantic Forest habitats. Out of the 35 OTUs, 24 were registered in Ubarana and 16 in Nazaré Paulista. Only OTU1, OTU2, OTU7, OTU10 and OTU20 occurred in both Nazaré Paulista and Ubarana (Supplemental Data 3). Average number of OTUs registered in Ubarana high-quality sites was 6.6 (ranging from 2 to 14 OTUs) and 6.25 for Ubarana low quality sites (ranging from 3 to 8 OTUs). While for Nazaré Paulista, average number of OTUs registered in high quality sites was 3.75 (ranging from 2 to 5 OTUs) and 1.8 for low quality sites (ranging from 1 to 4 OTUs).

Demultiplexed 454 sequence data for each of the 35 samples analysed herein are in NCBI’s Sequence Read Archive under project PRJNA645344 (https://www.ncbi.nlm.nih.gov/sra/).

### Alpha and beta diversity

The results of the model selection explaining richness were similar for both response variables (number of OTUs and phylogenetic diversity). In both cases, the model including only forest quality as fixed factor was selected as the best model, and the additive model including sampling month as second-best model (Supplemental Data 2 Table S1). While the additive model was equally plausible to the best model explaining the variation in *number of OTUs* (i.e., deltaAICc<2) it was not selected for *phylogenetic diversity*. The two models including forest quality thereby accumulated an AICc-weight of 94% (*number of OTUs*) 95% (*phylogenetic diversity*). Model estimates indicated that sampling sites within high-quality habitat had higher *number of OTUs* observed as well as higher *phylogenetic diversity* (Figure 2).

**Figure 2.**
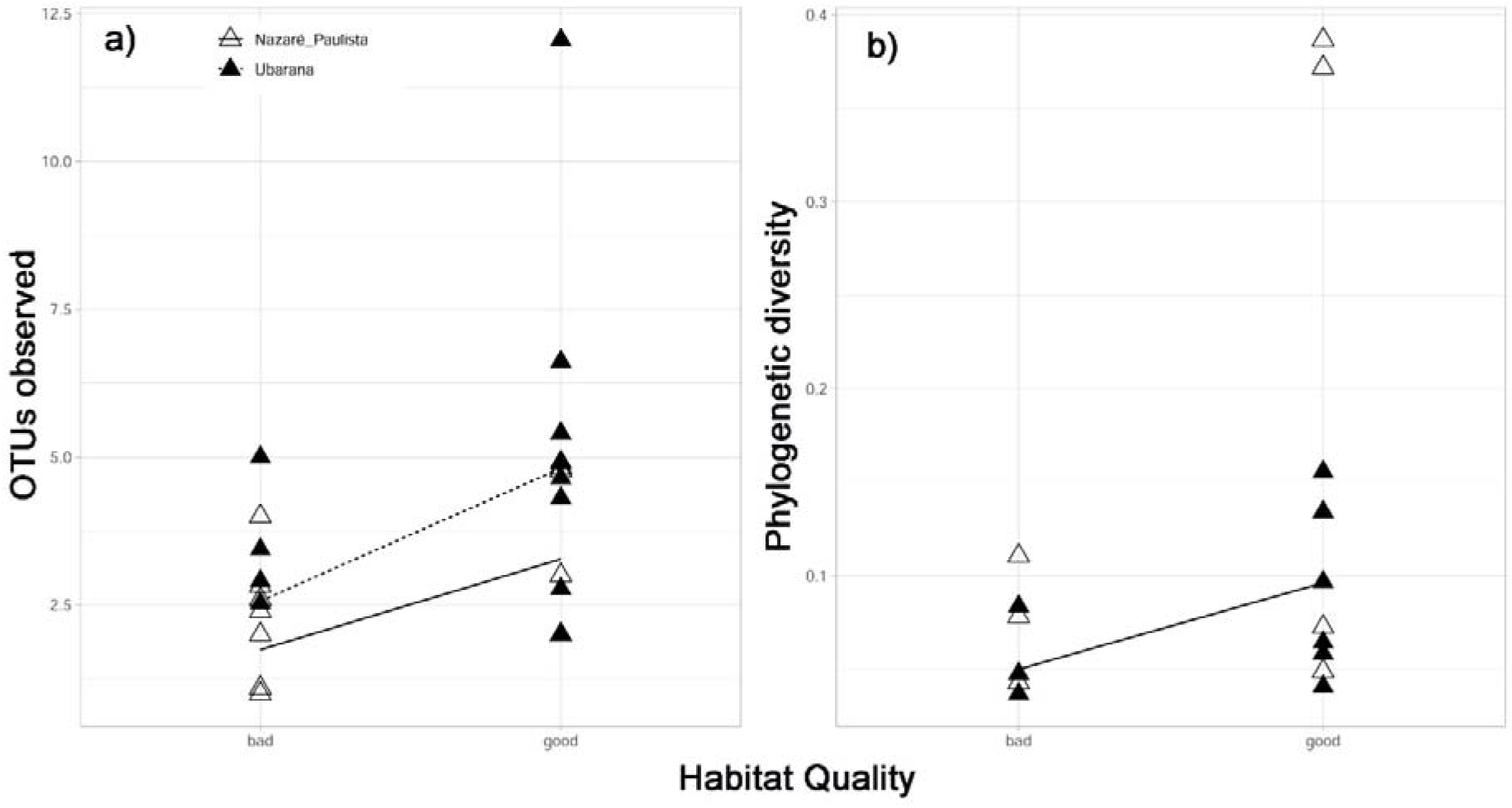
Predictions of best model explaining variation in number of observed OTUs (a) and phylogenetic diversity (b). Open triangles and smooth line: samples from Nazaré Paulista, filled triangles and dashed line: samples from Ubarana. Note that predictions for the increase in phylogenetic diversity are similar between the two regions, which led to superimposition of the respective lines in b).

Communities were significantly dissimilar among sampling areas (Supplemental Data 2). Additionally, most OTUs were highly specific to either high- or low-quality habitat, with only twelve OTUs occurring in both habitat classes (Supplemental Data 3), leading to a significant effect of habitat quality on community dissimilarity independently on the distance metric used (Supplemental Data 2 Table S2).

Accordingly, high- and low-quality sampling sites were clearly separated by NMDS (Figure 3).

**Figure 3.**
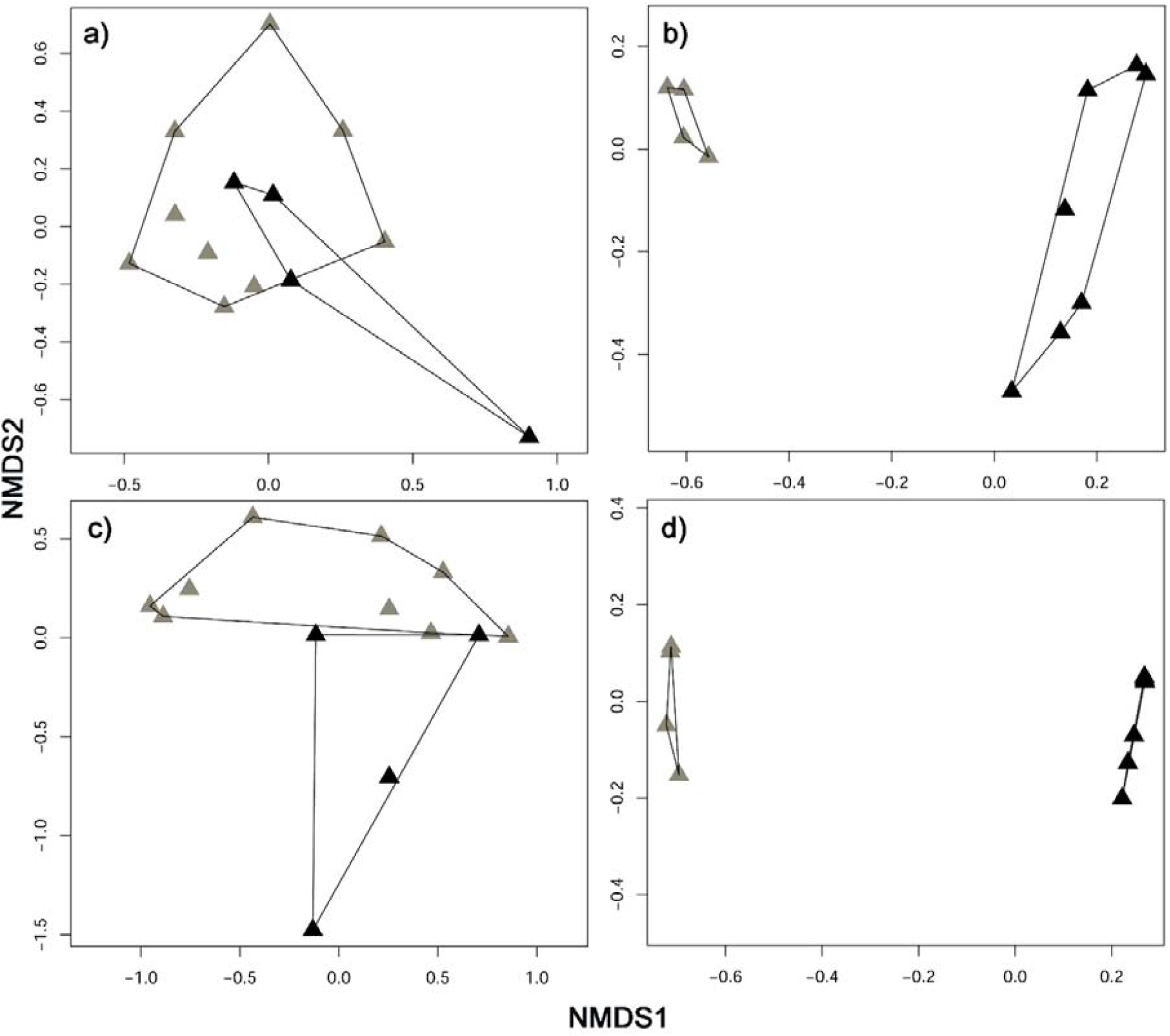
Non-metric multi-dimensional scaling (NMDS) of pairwise comparison of Collembola communities in Ubarana (a, b) and Nazaré Paulista (c, d) based on Jaccard- (a, c) and Bray-Curtis-dissimilarity (b, d). Grey symbols: sampling sites in high forest quality areas; black symbols: sampling sites sampling sites in low forest quality areas; Polygons enclose sites within same forest quality.

## Discussion

Metabarcoding represents a useful technique for capturing soil species-as well as community-level information, thereby providing useful insight on soil- and ecosystem-health. For example, Yang et al. (2014) found that leaf litter samples were considerably more informative in differentiating between habitat types than above-ground/aerial measures (Malaise trapping or fogging). Among soil invertebrates, Collembola are of special interest given their proven potential as ecological and biological indicators (e.g., Arenhardt et al., 2021; Cassagne et al., 2006). Therefore, by increasing the speed and accuracy of Collembola community analyses as well as decreasing costs, our primer set can provide considerable benefits for biomonitoring objectives.

### Potential for excluding non-target species

The D2 marker has previously been used in a variety of invertebrate molecular studies requiring species-level resolution (Deng et al., 2012; Schneider et al., 2011; Sonnenberg et al., 2007; Zhou et al., 2007). This domain has highly conserved flanking regions that can be used to anchor highly conserved primers across entire taxa. Our results show that the D2 segment of ribosomal DNA’s 28S operon can substantially facilitate the use of Collembola as an ecological indicator, as it provides efficient taxonomic resolution for these miniscule organisms, whose identification is often limited by their size.

The D2 primers herein allow for the exclusive amplification of Collembola from bulk samples with minimal PCR bias. The pipeline obviates time-consuming sorting and does not require that samples be morphologically preserved. It is especially appropriate when the sampling protocols used yield large and diverse amounts of bycatch (such as pitfall trapping), which, in previous Collembola metabarcoding studies, competed for large portions of the flow-cell yield (Saitoh, et al., 2016).

In highly polymorphic protein-coding loci (e.g., cytochrome oxidase I; COI) frequent synonymous mutations at third nucleotide sites make primer design either unstable or leads to primer bias because of polymorphisms (Clarke et al., 2014; Deagle et al., 2014; Pedro et al., 2020).

We explicitly show that, after sequence quality filtering, no non-target amplicons are derived. Moreover, the low degeneracy minimizes PCR bias and the primers’ species diagnostic performance is similar to both 16S and COI. Saitoh (2016) found a strong correlation between Collembola biomass and normalized sequence reads (R = 0.91–0.99), suggesting primers had little primer bias. However, in natural bulk samples, primers amplified a substantial proportion of non-targets (CO1 results produced 53% non-targets and 16S produced 35%), a substantial loss of flow cell capacity.

### Dearth of reference databases

The D2 marker may currently be inappropriate in contexts where species diagnosis is required (such as in pest detection or toxicology assays) because of the relatively small taxonomic reference databases. However, many soil biomonitoring studies, rely on sample comparisons with reference sites, rather than molecular taxonomy *per se* (e.g., comparisons between pristine forests and perturbed sampling sites (Russell and Alberti, 1998; Arboláez et al., 2023)). Moreover, although other markers, notably COI, have substantially more complete database, they are nonetheless probably still underrepresented because of the sheer number of species thought to still be described, especially in tropical soils (Bernard and Felderhoff, 2007; Cicconardi et al., 2013).

## Acknowledgments

This research received R&D funding from ANEEL (*Agência Nacional de Energia Elétrica*; grant number 0064-1035/2014) and also from *Projeto LIRA - Legado Integrado da Região Amazônica*, with funding partners *Fundo Amazônia/Banco Nacional de Desenvolvimento Econômico e Social* and the *Gordon and Betty Moore Foundation*. Sample collections were undertaken with permit number 54835 administered by the Brazilian Federal agency SISBIO. We would like to thank IdeaWild (https://ideawild.org) for important equipment contributions to this project.

## Notes

### Competing Interest Statement

The authors have declared no competing interest.

