## SupplementalData2 for "Ecological monitoring using Collembola metabarcoding with extremely low bycatch amplification"

Supplemental Data for: Collembola-based ecological monitoring through targeted metabarcoding with extremely low bycatch amplification.

Pitfall sampling protocol:

Invertebrate sampling was undertaken with “mini” pitfall traps improvised from 50-ml Falcon tubes half-filed with absolute ethanol and protected from rain and detritus with a plastic plate. The lip of the tube was fitted with an improvised inverted plastic funnel to limit ethanol evaporation and fitted with a 5-mm screen to prevent entry of taxa larger than ~2 mm. Three such pitfalls were used per sampling location, each placed within a radius of approximately 3 m.

We sampled in two regions of São Paulo State, Brazil. The first location, in Ubarana municipality (approximately 21°14’09” S, 49°43’12” W), consisted of a remnant central forest core adjacent to a hydroelectric reservoir. This was flanked by a narrower strip of forest at its south and a sparsely wooded strip at its north.

Four sampling points were located in a narrow strip of forest at its north characterized as LOW quality (because of its open canopy and a *Brachiaria*-dominated understory), five sampling points were located within the forest remnant, and four sampling points in the southern forest strip. Both the forest remnant and the southern forest strip were classified as HIGH quality (closed canopy, understory without grass).

At the second location, in Nazaré Paulista (23°12’48” S, 46°21’58” W) four sampling locations were located within HIGH-quality remnant forests, and 11 sampling locations within LOW-quality habitat (grasslands with sparse tree and shrub cover). Both sampling locations are characterized as Atlantic Forest *(http://mapas.sosma.org.br).* A google .kmz file is provided as Supplemental data 1 with specific coordinates. Unequal numbers of samples from high- and low-quality habitat was due to the loss of several samples destroyed by vandalism and wildlife. Pitfall traps were deployed for approximately 30 days beginning in September 2015 in Ubarana) and July 2016 in Nazaré Paulista (Supplemental data 1).

Alpha and beta diversity estimates:

Alpha diversity was estimated as the number of OTUs (MOTHUR command summary.single) and the scaled phylogenetic diversity (command phylo.diversity). In analyses involving phylogenetic diversity, we calculated inter-OTU distances based on a Neighbour-Joining tree (pairwise deletions and K2P-distance) created in MEGA (Kumar et al., 2016). In all diversity calculations, we used the MOTHUR command subsample=T to correct for disparate sequencing depth among the collections.

In order to assess the impact of forest quality on Collembola alpha-diversity, we used generalized linear mixed models (GLMMs). We constructed a candidate model set including a constant model (no fixed factors) as a reference, two simple models (with either quality or collection date as unique fixed factors, respectively), and one additive (including the combination of quality and month). In all candidate models we included “area” as a random factor, given that samples originated from two different areas. Both alpha diversity measures were modelled as lognormally distributed. For each of the two explanatory variables, we ran one model selection including the candidate model set. Models were compared using the Akaike Information Criterion adjusted for small sample sizes (AICc, Burnham & Anderson, 2002).

We calculated two beta-diversity metrics of pair-wise community dissimilarity with the vegan R package (Oksanen et al., 2022): the Jaccard index (occurrence-based index) and the Bray-Curtis dissimilarity index (abundance-based). Non-metric multidimensional scaling (NMDS) was used for ordination plots based on each distance matrix. Permutational multivariate analysis of variance (PERMANOVA; (Anderson, 2006) was used to test for effects of forest quality on beta-diversity. Separate PERMANOVAs for each sampling location were done and models included, in order of addition: “sampling region”, “sampling month”, and “forest quality” as factors. F- and p-values were calculated using 10000 permutations.

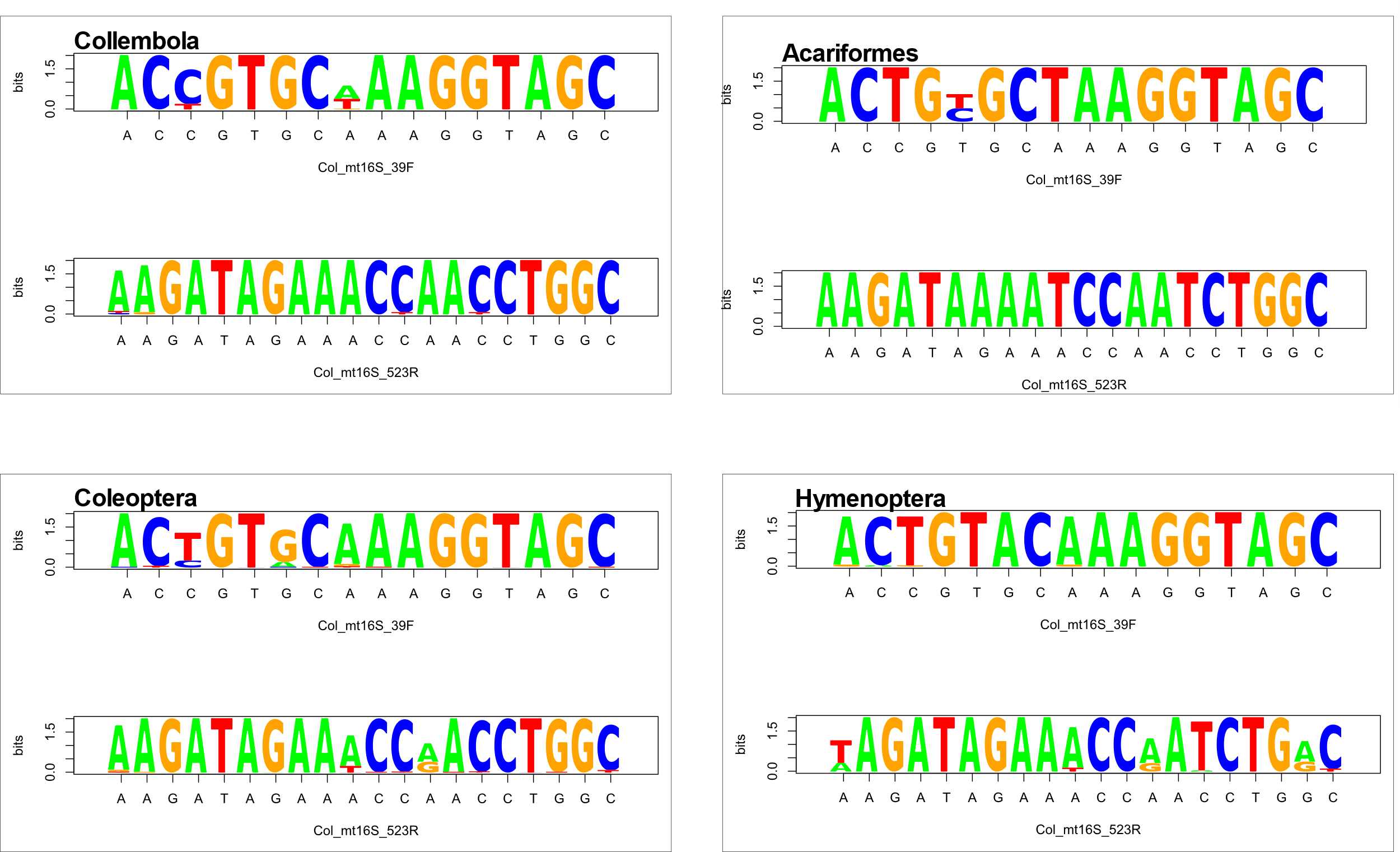

Figure S1.

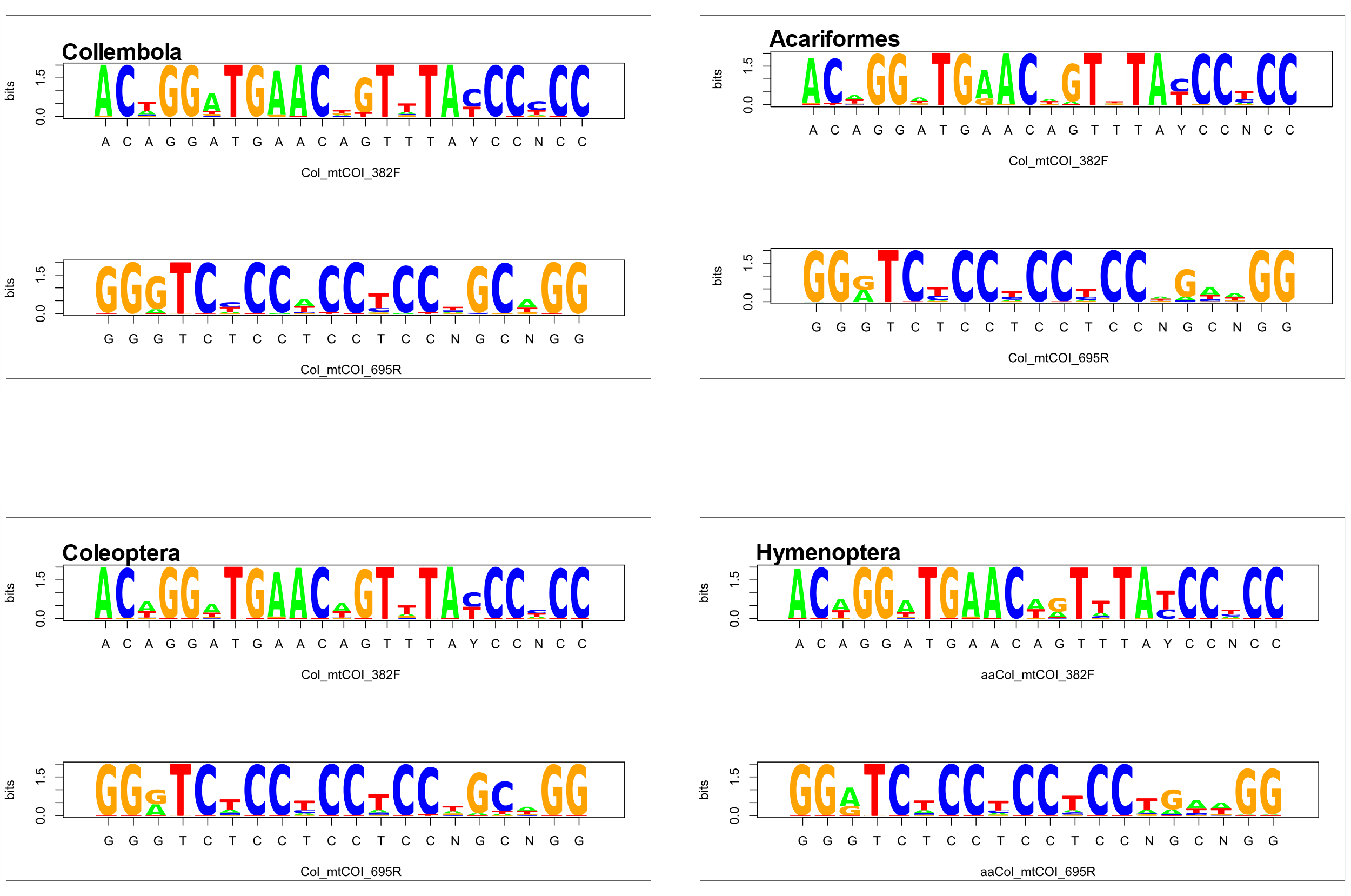

Figure S2.

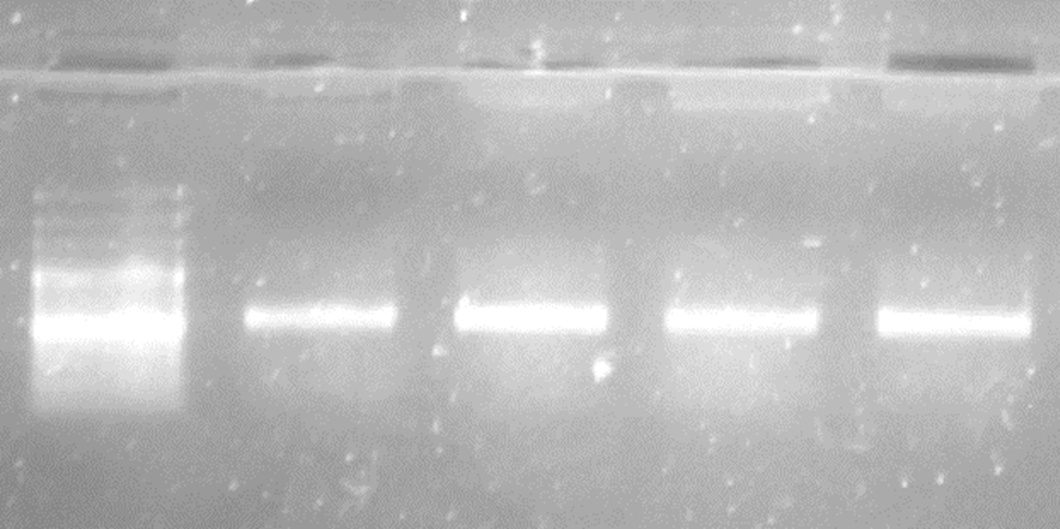

Figure S3. Product of first of the two-step PCR using the CollembolaF and CollembolaR primers with Illumina adaptors. Viewed on 1.5% agarose gel stained with ethidium bromide.

Table S1. Results of model selection for a) number of observed taxonomic units (OTUs), and b) phylogenetic diversity. Model: fixed factor(s) included in the model (constant = null model including only the intercept). K: number of parameters; AICc: Akaikes Information Criterion adjusted for small samples size; Delta_AICc: difference in AICc value compared to the best model; AICcWt: Akaike weigth; Cum.Wt: cumulative Akaike weights; LL: log-likelihood. All models included “sampling region” as a random factor.

| a) OTUs |  |  |  |  |  |  |
| --- | --- | --- | --- | --- | --- | --- |
| Model | K | AICc | Delta_AICc | AICcWt | Cum.Wt | LL |
| **quality** | **4** | **50.84** | **0.00** | **0.68** | **0.68** | **-20.55** |
| **quality + month** | **5** | **52.72** | **1.88** | **0.27** | **0.94** | **-19.99** |
| constant | 3 | 56.58 | 5.74 | 0.04 | 0.98 | -24.79 |
| month | 4 | 58.26 | 7.42 | 0.02 | 1.00 | -24.26 |
| b) Phylogenetic diversity | |  |  |  |  |  |
| Model | K | AICc | Delta_AICc | AICcWt | Cum.Wt | LL |
| **quality** | **4** | **53.88** | **0.00** | **0.78** | **0.78** | **-22.07** |
| quality + month | 5 | 56.83 | 2.95 | 0.18 | 0.95 | -22.05 |
| constant | 3 | 60.10 | 6.23 | 0.03 | 0.99 | -26.55 |
| month | 4 | 62.17 | 8.30 | 0.01 | 1.00 | -26.22 |

Table S2. Results of PERMANOVA based on two beta-diversity indices including both sampling areas (Ubarana and Nazaré Paulista) showing effects of fixed factors (sampling region, sampling month and forest quality) on pairwise community dissimilarity. Df: degrees of freedom; SumsOfSqs: sums of squares; MeanSqs: mean sums of squares; F.Model: pseudo-F statistic; R2: percent variance explained; Pr(>F): probability of finding F-statistic as large or larger assuming null hypothesis of no effect. Significant p-values are in bold.

|  | Df | SumsOfSqs | MeanSqs | F.Model | R^2^ | Pr(>F) |
| --- | --- | --- | --- | --- | --- | --- |
| a) Jaccard index | |  |  |  |  |  |
| area | 1 | 3.11 | 3.11 | 13.33 | 0.30 | **0.0001** |
| month | 1 | 0.44 | 0.44 | 1.88 | 0.04 | 0.0716 |
| quality | 1 | 1.36 | 1.36 | 5.81 | 0.13 | **0.0001** |
| Residuals | 24 | 5.60 | 0.23 |  | 0.53 |  |
| Total | 27 | 10.50 |  |  | 1.00 |  |
| b) Bray-Curtis dissimilarity | | |  |  |  |  |
| area | 1 | 3.55 | 3.55 | 19.05 | 0.35 | **0.0001** |
| month | 1 | 0.30 | 0.30 | 1.61 | 0.03 | 0.1548 |
| quality | 1 | 1.78 | 1.78 | 9.54 | 0.18 | **0.0001** |
| Residuals | 24 | 4.48 | 0.19 |  | 0.44 |  |
| Total | 27 | 10.11 |  |  | 1.00 |  |

Anderson, M. J. (2006). Distance-based tests for homogeneity of multivariate dispersions. *Biometrics*, *62*(1), 245–253. https://doi.org/10.1111/j.1541-0420.2005.00440.x

Burnham, K. P., & Anderson, D. R. (2002). Model selection and multimodel inference - a practical information-theoretic approach. In *Model Selection and Multimodel Inference* (2nd ed.). Springer New York. https://doi.org/10.1007/b97636

Kumar, S., Stecher, G., & Tamura, K. (2016). MEGA7: Molecular Evolutionary Genetics Analysis Version 7.0 for Bigger Datasets. *Molecular Biology and Evolution*, *33*(7), 1870–1874. https://doi.org/10.1093/molbev/msw054

Oksanen, J., Simpson, G. L., & Blanchet, F. G. (2022). *Vegan: Community Ecology Package. R package version 2.6–2.* (p. 295). https://github.com/vegandevs/vegan NeedsCompilation
